# Discovering novel long non-coding RNA predictors of anticancer drug sensitivity beyond protein-coding genes

**DOI:** 10.1101/666156

**Authors:** Aritro Nath, Eunice Y.T. Lau, Adam M. Lee, Paul Geeleher, William C.S. Cho, R. Stephanie Huang

## Abstract

Large-scale cancer cell line screens have identified thousands of protein-coding genes (PCGs) as biomarkers of anticancer drug response. However, systematic evaluation of long non-coding RNAs (lncRNAs) as pharmacogenomic biomarkers has so far proven challenging. Here, we study the contribution of lncRNAs as drug response predictors beyond spurious associations driven by correlations with proximal PCGs, tissue-lineage or established biomarkers. We show that, as a whole, the lncRNA transcriptome is equally potent as the PCG transcriptome at predicting response to hundreds of anticancer drugs. Analysis of individual lncRNAs transcripts associated with drug response reveals nearly half of the significant associations are in fact attributable to proximal *cis*-PCGs. However, adjusting for effects of *cis-*PCGs revealed significant lncRNAs that augment drug response predictions for most drugs, including those with well-established clinical biomarkers. In addition, we identify lncRNA-specific somatic alterations associated with drug response by adopting a statistical approach to determine lncRNAs carrying somatic mutations that undergo positive selection in cancer cells. Lastly, we experimentally demonstrate that two novel lncRNA, *EGFR-AS1* and *MIR205HG*, are functionally relevant predictors of anti-EGFR drug response.

## Background

LncRNAs are transcripts greater than 200 nucleotides in length that do not contain protein-coding sequences. They act as key regulators of gene expression (Mattick and Rinn 2015), controlling a diverse set of transcriptional and post-transcriptional processes, including chromatin remodeling, RNA splicing and transport, and protein synthesis (Wang and Chang 2011; Kopp and Mendell 2018). Although, less than 1% of lncRNAs have been functionally characterized (Quek et al. 2015), comprehensive characterization of lncRNA across thousands of tumors suggest pervasive dysregulation of the lncRNA transcriptome at rates similar to protein-coding genes (PCGs) (Iyer et al. 2015; Yan et al. 2015). In addition, some lncRNAs function as either oncogenes or tumor suppressor genes in human cancers (Huarte 2015; Schmitt and Chang 2016).

Several large-scale cancer cell line screens systematically investigated the response to hundreds of drugs to identify genomic and transcriptomic biomarkers of cancer drug response (Barretina et al. 2012; Garnett et al. 2012; Basu et al. 2013; Seashore-Ludlow et al. 2015; Iorio et al. 2016b). These studies expanded the repertoire of somatic alterations and gene expression biomarkers linked with drug response but focused exclusively on PCGs. Considering less than 2% of the genome codes for PCGs (Djebali et al. 2012) with nearly 70% of the genome transcribed into non-coding RNAs (Derrien et al. 2012), it seems that the mechanisms of anticancer drug response cannot be explained by PCGs alone (Malek et al. 2014). Subsequently, a recent study reported lncRNA models are better predictors of drug response compared to PCGs for several drugs (Wang et al. 2018). However, lncRNAs are expressed with a high degree of tissue-specificity and the expression of genic lncRNAs tends to be strongly correlated with the expression of PCGs on complementary strands (Derrien et al. 2012). Therefore, the identification of novel lncRNA biomarkers associated with anticancer drug response requires careful consideration of the potential confounding influence of the tissue lineage along PCGs proximal to the lncRNAs.

Here we report the results of a systematic investigation of the lncRNA transcriptome and genome of cancer cell lines and large-scale drug screens to establish a pharmacogenomic landscape of lncRNAs. We use regularized regression models to predict drug response using lncRNA transcriptome to demonstrate its potency compared to PCGs. To guide the discovery of individual lncRNA biomarkers, we delineate the effects of *cis*-PCGs on drug-lncRNA associations in regression models. In addition, we identify lncRNA-specific somatic mutations undergoing positive selection in cancer cells and determine their associations with drug response. We further investigate the contribution of lncRNAs in predicting the response for drugs with clinically actionable PCG biomarkers. Based on our analysis, we highlight the role of *EGFR-AS1* and *MIR205HG* as predictors of anti-EGFR therapeutic response independent from *EGFR* somatic mutations and experimental confirm their potential as erlotinib-response biomarkers in lung cancer cells.

## Results

Despite the tremendous success of cancer cell line screens in discovering novel PCG biomarkers of drug response, the contribution of lncRNAs in cancer pharmacogenomics is poorly established. To systematically determine the relevance of lncRNAs as anticancer drug response biomarkers, we propose the following framework to delineate their contribution as response predictors while accounting for the effects of proximal *cis*-PCGs (within ±500kb) and known biomarkers (Figures 1A-D, Supplementary Figures 1A, B). In addition, we determine the contribution of somatic alterations specific to lncRNAs in drug response using a statistical approach to determine positively selected mutations (Figure 1E). Finally, we experimentally validate the functional role of two novel lncRNA predictors of anti-EGFR drug response (Figure 1F).

**Figure 1:**
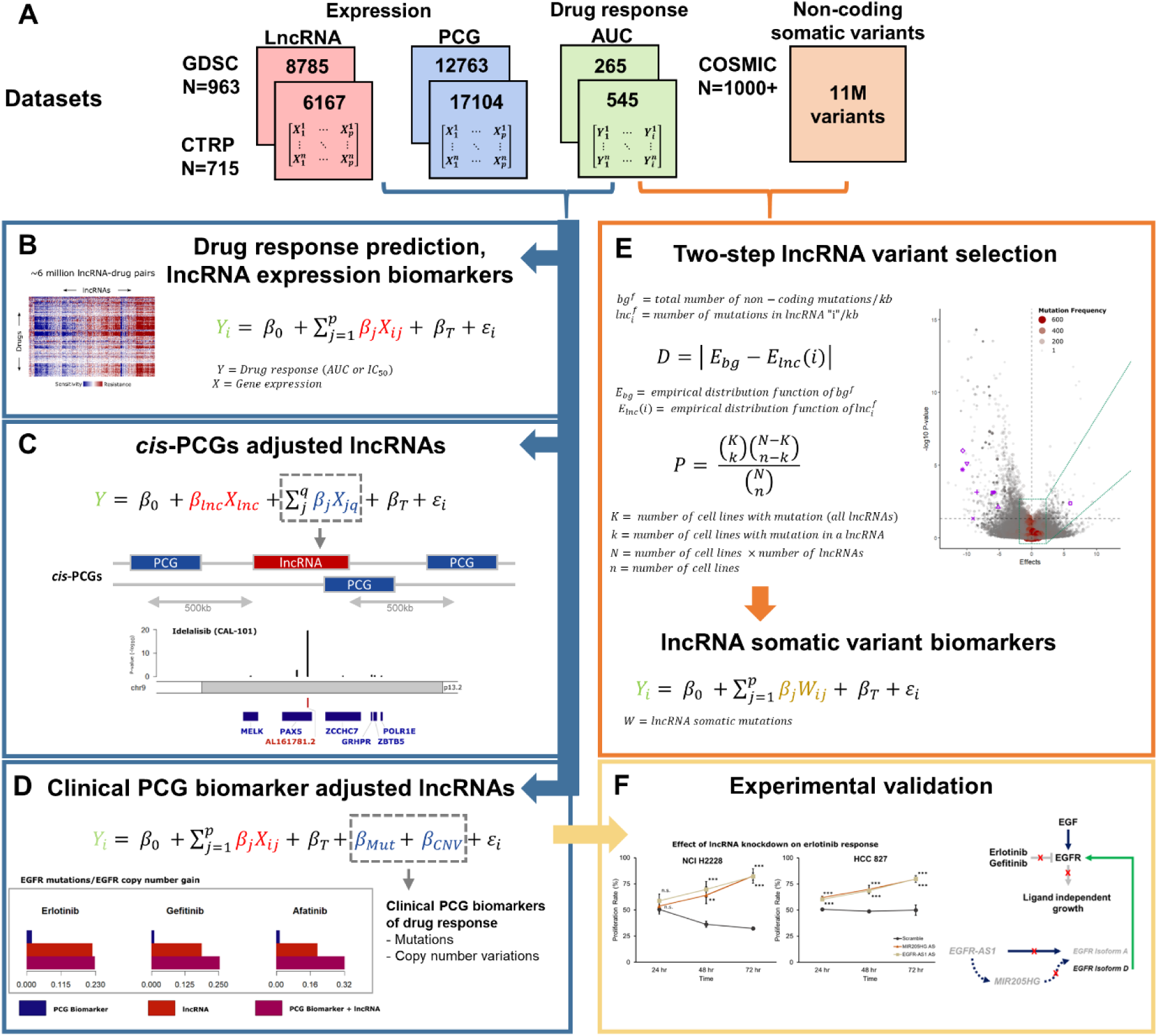
Framework for novel lncRNA biomarker discovery. **A.** Datasets used in the study, including gene expression (PCG, lncRNA) and drug response profiles corresponding to Genomics of Drug Sensitivity in Cancer (GDSC) and Cancer Therapeutics Response Portal (CTRP) cell lines, and non-coding somatic variants from Catalogue of Somatic Mutations in Cancer (COSMIC). “N” indicates the number of cell lines in each dataset, while the number of lncRNA, PCGs or drugs with the area under the curve (AUC) of drug response in each dataset are indicated inside the colored boxes. **B.** Linear model for predicting drug response (AUC) using the PCG or lncRNA transcriptome **C.** Determining significance drug:lncRNA associations after adjusting for the expression levels of neighboring *cis*-PCGs within a ±500kb window. **D.** Identification of significant drug:lncRNA associations after adjusting for the mutation or copy number variation status of clinically-established PCG biomarkers of drug response. **E.** A two-step statistical approach to determine lncRNAs with somatic mutations that undergo positive selection in cancer cell lines. **F.** Experimental validation of candidate lncRNAs that augment clinical biomarkers of drug response

### Determining the contribution of lncRNA transcriptome as a predictor of anticancer drug response

An important finding reported by the cancer cell line screens was the ability to implement machine-learning algorithms that accurately predicted drug response using the baseline PCG transcriptome of cancer cells. Thus, we first compared the ability of lncRNA transcriptome to predict response to 265 and 545 compounds from the GDSC and CTRP screens respectively. In both screens, the lncRNA transcriptome was equally potent at predicting response to individual drugs as PCGs (CTRP Spearman’s ρ = 0.93; GDSC Spearman’s ρ = 0.98) (Figure 2A), with no difference in median prediction accuracies across all drugs (CTRP P = 0.17; GDSC P = 0.32) (Supplementary Figure 1C). The drug GSK-J4, a potent and highly selective inhibitor of H3K27 histone demethylases JMJD3 and UTX, was an exception that was predicted with better accuracy using lncRNA transcriptome. For the set of drugs that were common to both screens, consistent prediction accuracies were achieved using both lncRNA (P = 0.24) and PCG models (P = 0.07) (Supplementary Figures 1D and 1E).

**Figure 2:**
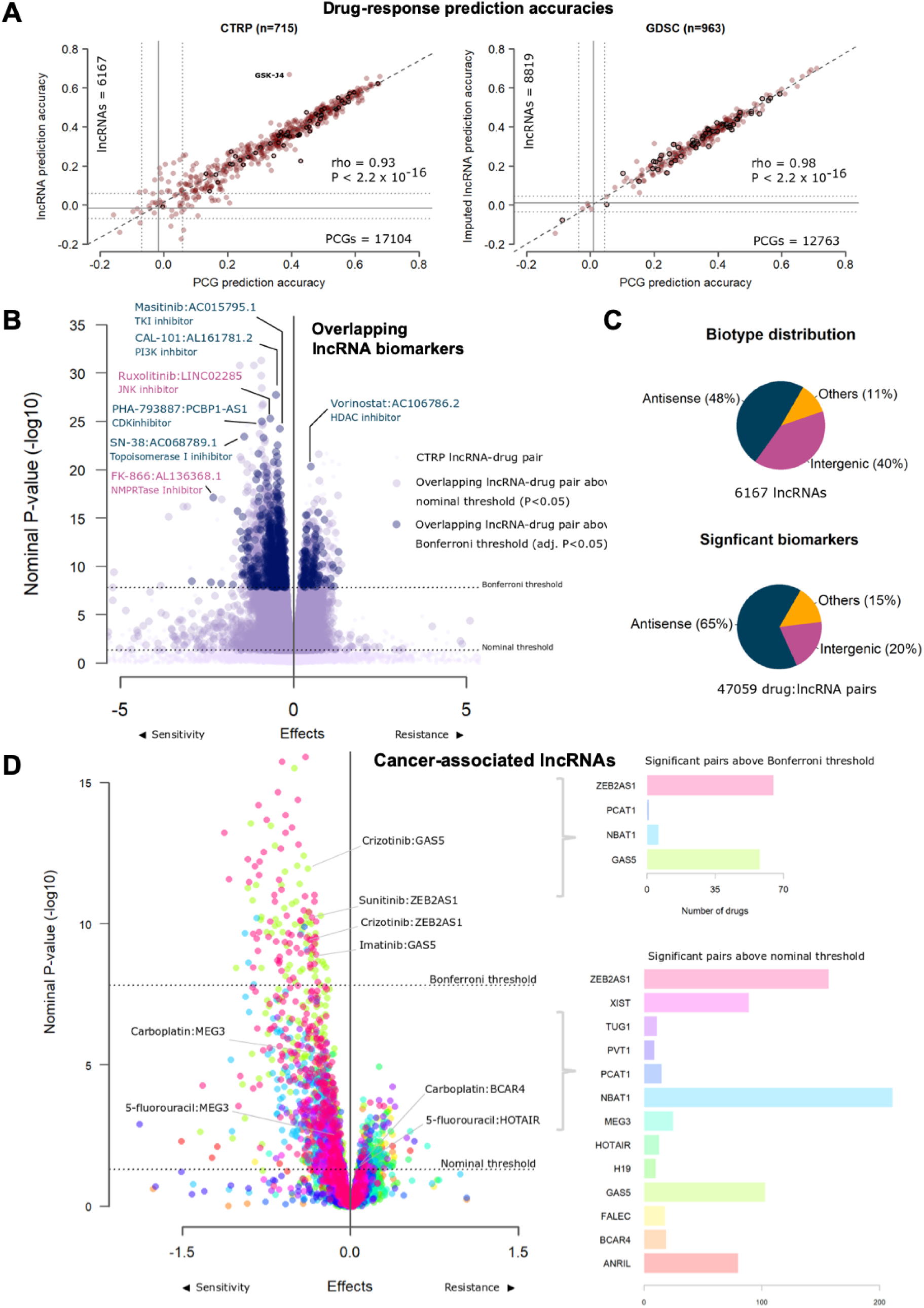
LncRNAs as predictors and biomarkers of anticancer drug response. **A.** Scatterplot of 545 CTRP (C) or 265 GDSC (D) prediction accuracies using PCG transcriptome (X-axis) or imputed lncRNA transcriptome (Y-axis) as predictors. Each point on the scatter plot shows the accuracy of predicting response to a drug by using models generated using PCG or lncRNA transcriptome. The bolded points are drugs common to both GDSC and CTRP. Grey lines indicate prediction accuracies with confidence intervals from a null model. **B.** Volcano plots of drug-lncRNA associations in the cohort of drug-lncRNA pairs common to CTRP and GDSC screens displaying effect sizes (X-axis) and P-values (Y-axis) of the regression analyses. The light-blue enlarged circles indicate significant (FWER adjusted) CTRP drug-lncRNA pairs also significant in GDSC at the nominal threshold, while dark-blue enlarged circles are significant in GDSC at the FWER threshold. **C.** Distribution of lncRNA biotypes across all cell lines and within the subset of significant drug-lncRNA pairs (FWER adjusted) identified from linear regression analysis of the CTRP screen adjusted for tissue-type. **D.** Volcano plot displaying effect sizes (X-axis) and P-values (Y-axis) of oncogenic and tumor suppressor lncRNAs associated with drug response after adjusting for *cis*-PCGs. The bar plots on the side indicate the frequency of drugs associated with the lncRNAs at the nominal threshold and FWER-adjusted threshold.

Next, we modeled drug response as a function of individual lncRNA transcripts to determine significant biomarkers. Across the set of drug-lncRNA pairs common to both CTRP and GDSC screens, 68% of the significant (FWER < 0.05) CTRP drug-lncRNA associations were also significant in GDSC at the nominal threshold (P < 0.05), while about 28% were significant at the FWER threshold (Figure 2B). Both antisense and intergenic transcripts were represented in the cohort of top significant overlapping drug pairs; for example, vorinostat resistance was associated with the antisense lncRNA AC106786.2 or ruxolitinib sensitivity with the intergenic lncRNA LINC02285 (Figure 2B). The biotype of lncRNAs were nearly equally distributed between genic (including antisense) and intergenic RNAs, while antisense transcripts were over-represented in the cohort of significant (family-wise error rate [FWER] < 0.05) drug-lncRNA pairs as compared to intergenic transcripts (Tukey’s P < 10^−11^) (Figure 2C, Supplementary Table 1).

Next, we evaluated the pharmacogenomic relevance of the well-characterized lncRNAs have been implicated as oncogenes or tumors suppressors in human cancers (Gutschner and Diederichs 2014) (Figure 2D, Supplementary Figure 1C). For example, the expression of putative tumor suppressor lncRNA *MEG3* was associated with sensitivity to carboplatin and 5-fluorouracil, while the putative oncogenes *BCAR4* and *HOTAIR* were associated with resistance to carboplatin and 5-fluorouracil, respectively (Supplementary Table 3). These observations are in line with previous *in vitro* studies that show elevated *MEG3* expression to be associated with sensitivity (Cheng et al. 2015; Li et al. 2017a) while high *BCAR4* (Godinho et al. 2010) and *HOTAIR* (Kalinichenko et al. 2013) expression were linked with resistance to cytotoxic anticancer agents. Within in the cohort of significant cancer-associated lncRNAs, we observed the expression of *GAS5* and *ZEB2AS1* were associated with the sensitivity of more than 50 drugs (Figure 2D), suggesting these lncRNAs could be candidates for further evaluation as multi-drug response predictors. These results suggest the potential of known cancer-associated lncRNAs to serve as determinants of anticancer drug response.

### Characterizing the impact of proximal *cis*-PCGs on drug-lncRNA associations

The strong correlation between the expression levels of antisense lncRNAs and overlapping PCGs could potentially result in spurious associations with drug response. To deconvolute the contribution of lncRNAs from the effects of proximal PCGs, we next analyzed drug response as a function of lncRNA expression while adjusting for the effects of proximal *cis*-PCGs (any PCG located in the −500Kb region upstream of transcription start site to +500Kb downstream of transcript end). Nearly 50% of the remaining significant drug-lncRNA associations were affected by the expression levels of the *cis*-PCGs (Figure 3A). As expected, adjusting for *cis*-PCGs mostly affected the proportion of significant antisense lncRNAs over intergenic transcripts (Figure 3A).

**Figure 3:**
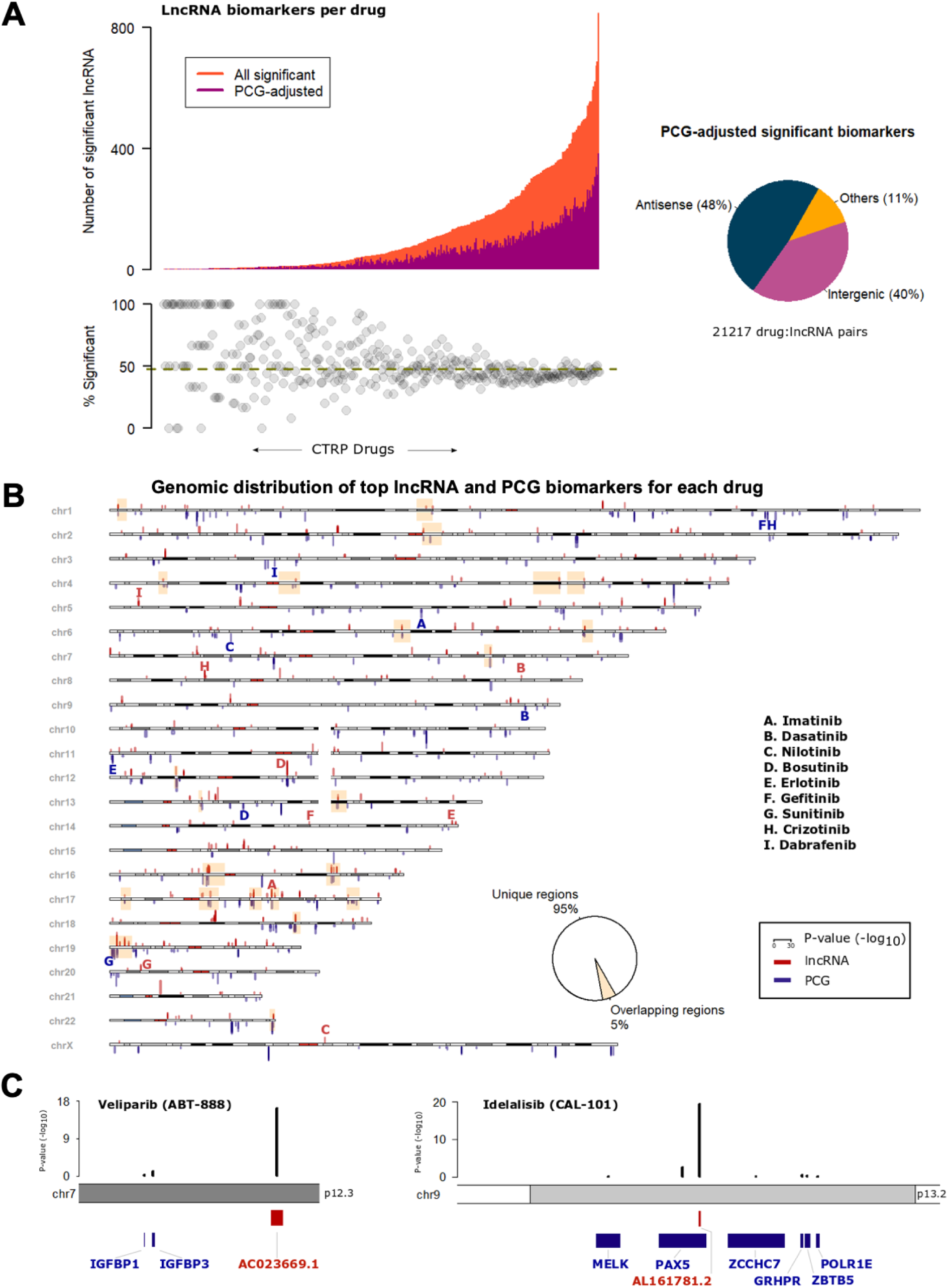
Relevance of cis-PCG adjusted lncRNA biomarkers. **A.** Distribution of significant lncRNAs associated with each CTRP drug (X-axis). The orange bars indicate a number of significant lncRNAs for a drug while the magenta bars indicate the number of lncRNAs that are statistically significant after adjusting for the expression levels of every *cis*-PCG (within 1 Mb) of the lncRNA transcript. Biotype distribution of the lncRNAs in the cohort of significant drug-lncRNA pairs adjusted for *cis*-PCGs. **B.** Ideogram of the human chromosomes displaying the top lncRNA and PCG associated with each CTRP drug. P-values of the drug-gene associations are indicated as red bars for lncRNAs above the chromosomes and as blue bars for PCGs below the chromosomes at their respective locus. The yellow highlighted boxes indicate the cytobands with overlapping top lncRNA and PCG loci associated with a drug. Bolded letters (red = lncRNA, blue = PCG) indicate signals associated with the set of drugs with clinically actionable biomarkers. **C.** Examples of top lncRNA associated with veliparib and idelalisib response, with black bars indicating adjusted P-values for lncRNA and *cis*-PCGs.

Given the obvious influence of proximal PCGs on drug-lncRNA associations and to better understand the genome-wide distribution of lncRNA biomarkers relative to PCGs, we mapped the significant lncRNA and PCG biomarker for all drugs analyzed in the CTRP screen (Supplementary Figures 2A and 2B). For individual drug, we also mapped the top predictive lncRNA or PCG biomarkers based on their genomic loci (Figure 3B). Interestingly, we observed that for most drugs, the top lncRNAs and PCG biomarkers were located on distinct loci. In fact, the top lncRNA and PCG loci overlapped for only 5% of the drugs analyzed. As a case in point, we highlight the set of drugs with established clinically actionable biomarkers (Relling and Evans 2015). For example, the top PCG markers for gefitinib and crizotinib response were located on chromosomes 1 while the lncRNAs markers were located on 14 and 8, respectively. Based on these observations, we propose that despite the obvious impact of proximal *cis-*PCGs, it is possible to discover strong drug-lncRNA associations by carefully accounting for the effects of *cis-*PCGs.

As additional examples, we zoom-in to highlight the top lncRNA associated with veliparib and idelalisib response along with the proximal *cis*-PCGs. The intergenic lncRNA *AC023669.1* was associated with veliparib response (P = 3.9 × 10^−17^) after adjusting for the neighboring PCGs *IGFBP1* (P = 0.7) and *IGFBP3* (P = 0.08) (Figure 3C). The antisense lncRNA *AL161781.2* was associated with idelalisib sensitivity (P = 2.9 × 10^−20^) while the expression levels of proximal *cis*-PCGs showed considerably weaker association with the drug (e.g. *PAX5* P = 0.002) (Figure 3C).

### LncRNAs augment drug response predictions from known PCGs biomarkers

Currently, a small number of PCG mutations and copy number variations (CNVs) are being used in the clinic as biomarkers to guide treatment decisions (Relling and Evans 2015). We evaluated if the top lncRNA biomarkers for such drugs provide any additional benefit over the clinical biomarkers. We modeled drug response as a function of lncRNA expression and known PCG biomarkers and compare the individual and combined contribution of each predictor at explaining the variability in drug response (Figure 4A). In the case of BCR-ABL targeting tyrosine kinase inhibitors (TKIs), like imatinib and nilotinib, the *BCR-ABL1* fusion event was a stronger predictor compared to the top lncRNA. However, the response to dasatinib, a TKI with several other targets besides BCR-ABL, was better explained by the lncRNA. Similarly, *BRAF* mutations were strong predictors of response to dabrafenib and trametinib; and *ERBB2* mutations/CNVs for lapatinib sensitivity. In each of these cases, the addition of the lncRNA biomarker improved the proportion of variance explained by the model compared to the PCG biomarker alone. Other mutations and CNVs in genes like *KIT*, *PDGFR*, *KRAS*, *ALK*, and *VHL* actually explained a very small proportion of the variance of the respective drugs and were supplemented by the addition of lncRNA biomarkers. Interestingly, *EGFR* mutations and CNVs, one of the most prominent examples of cancer pharmacogenomic biomarkers for the EGFR targeting TKIs, were also augmented by the inclusion of expression of the top lncRNA biomarker. These results provide strong evidence to support the utility of lncRNA expression as biomarkers for anticancer drugs beyond PCGs.

**Figure 4:**
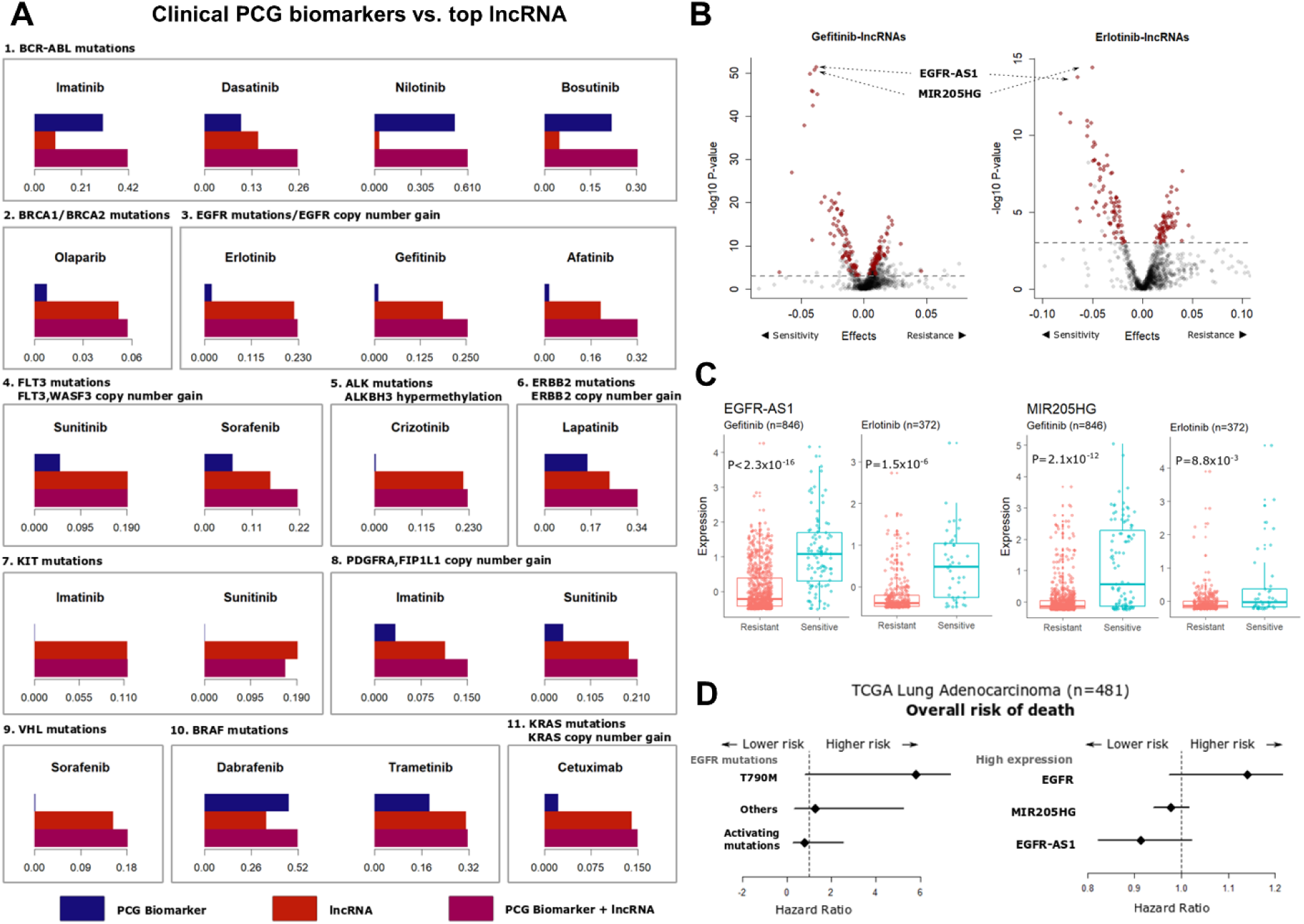
LncRNAs augment clinical drug response biomarkers. **A.** Barplots showing the proportion of variance in drug response (R^2^ of the regression model adjusted for tissue type) explained by the known PCG biomarker (blue bar), top lncRNA biomarker (red bar) or model combining the two (violet bar). The PCG biomarkers are listed above each sub-panel. **B.** Volcano plots showing lncRNAs associated with gefitinib or erlotinib response in the GDSC dataset adjusting for mutations and copy number variations in the *EGFR* gene. Each point on the plot represents a lncRNA-drug pair, with red points indicating lncRNAs common to both erlotinib and gefitinib above the nominal significance threshold of FDR < 0.05 (dashed grey line). **C.** Distribution of *EGFR-AS1* (top panels) and *MIR205HG* (bottom panels) expression in gefitinib or erlotinib resistant (AUC > 0.9) or sensitive (AUC < 0.9) cell lines. **D.** Hazard ratios obtained from Cox-proportional hazard models for overall survival of TCGA LUAD patients based on *EGFR* mutation status (top panel) or elevated expression levels of the *EGFR*, *EGFR-AS1* or *MIR205HG* genes.

We further evaluated the top lncRNA predictors of EGFR-targeting TKI response, as the inclusion of these lncRNAs in the model resulted in substantial improvement in the proportion of variability in drug response explained by EGFR mutations or CNVs alone. Somatic mutations in the *EGFR* tyrosine kinase domain, including in-frame deletions in exon 19, single nucleotide variations in exon 21 and amplification improve sensitivity, while an exon 20 (T790M) secondary mutation causes resistance to the anti-EGFR drugs gefitinib and erlotinib (Paez 2004; Pao et al. 2004; Liu et al. 2005; Moroni et al. 2005). However, these well-defined biomarkers can only explain a small proportion of the variance in drug response in (Figure 4A). This observation is consistent with data from non-small cell lung cancer (NSCLC) patients, where the response to anti-EGFR therapy is determined by EGFR activating mutations and CNVs in about 10-30% of patients (Gazdar 2009). However, about 1 in 4 patients that respond to gefitinib or erlotinib do not carry these activating alterations (Sharma et al. 2007). We found two lncRNAs, the EGFR antisense RNA 1 (*EGFR-AS1*; ENSG00000224057) and the MIR205 host gene (*MIR205HG*; ENSG00000230937), as the top two candidates biomarkers of anti-EGFR drug response independent of known PCG biomarkers (Figure 4B). The addition of *EGFR-AS1* and *MIR205HG* expression substantially improved the proportion of variance explained by the drug response models to 12-18% for erlotinib and 25-30% for gefitinib when combined with the EGFR functional events across all cell lines (Supplementary Figure 3A). Without considering EGFR functional events, both *EGFR-AS1* and *MIR205HG* expression levels were higher in the cells sensitive to gefitinib or erlotinib (Figure 4C, Supplementary Figure 3B). Moreover, the drug-lncRNA associations for *EGFR-AS1* (erlotinib P = 1.37×10^−10^; gefitinib P = 2.2×10^−16^) and *MIR205HG* (erlotinib P = 2.04×10^−4^; gefitinib P = 2.2×10^−16^) were significant after adjusting for *EGFR* mutations and CNVs (Supplementary Figure 3C).

Building on these results, we validated the correlation between *EGFR-AS1* and *MIR205HG* expression with imputed erlotinib response in lung adenocarcinoma (LUAD) patients from the cancer genome atlas lung adenocarcinoma (TCGA) project (Geeleher et al. 2017). The analysis of imputed drug response showed higher expression of both *EGFR-AS1* and *MIR205HG* were associated with sensitivity to erlotinib (Supplementary Figures 4A – D). The significance of correlation between the lncRNAs and erlotinib response was within the same order of magnitude as *EGFR* mutation status (Supplementary Figures 4E, F).

We next evaluated the impact of the candidate lncRNA biomarkers for on patient survival outcomes, in comparison with EGFR mutation status. As expected in the TCGA LUAD cohort, the presence of *EGFR* secondary resistance mutation (T790M) or high *EGFR* expression were both associated with worse overall prognosis (Figure 4D). In contrast, the elevated expression of *MIR205HG* and *EGFR-AS1* were associated with reduced risk of lung cancer death, a trend similar to the presence of EGFR activating mutations (Figure 4D). These results are consistent in the directionality suggested by our analysis, that is, the expression of both *EGFR-AS1* and *MIR205HG* are indicative of improved sensitivity and better prognosis. Thus, further inquiry into the prognostic value of *MIR205HG* and *EGFR-AS1* for anti-EGFR therapeutics will be crucial.

### Determining lncRNA-specific genomic alterations associated with drug response

The majority of the mutation profiles utilized in existing cancer pharmacogenomic studies were generated using exome sequencing. As a result, the impact of somatic variants that specifically affect the lncRNA genome has largely remained unexplored. The study of lncRNA variants is complicated by the lack of a clear definition of ‘passenger’ and ‘driver’ non-coding mutations. Two recent efforts attempted to identify driver lncRNA mutations that were positively selected in human cancers (Juul et al. 2017; Lanzós et al. 2017). In some form, each method identified non-coding loci or non-coding genes that were mutated at a frequency significantly greater than the background rate across all non-coding loci or across samples, respectively. We adopted a similar framework to define positively selected lncRNA-specific somatic variants using whole genome sequencing data from about 1000 catalog of somatic mutations in cancer (COSMIC) cell lines (Iorio et al. 2016b). We excluded all genic non-coding variants (intron, promoter or untranslated regions of PCGs) and identified lncRNA genes with mutation frequencies greater than the length-adjusted background non-coding mutation frequency.

We analyzed drug response as a function of the mutation status of each candidate lncRNA (Figure 5A, Supplementary Figures 5A, B). In contrast with lncRNA expression, only a small set of lncRNA mutations undergoing positive selection were associated with drug response (Inset Figure 5A, Supplementary Figures 5C, 5D). In the case of drugs with actionable PCG biomarkers, we found drugs with a similar mechanism of actions tend to share the top lncRNA mutation predictors (Figure 5A). For example, *AC007405.1* (ENSG00000234350) was the top lncRNA associated with sensitivity to both erlotinib and gefitinib (Figure 5B), suggesting a possible functional link between these lncRNA loci and the mechanism of action of the drugs.

**Figure 5:**
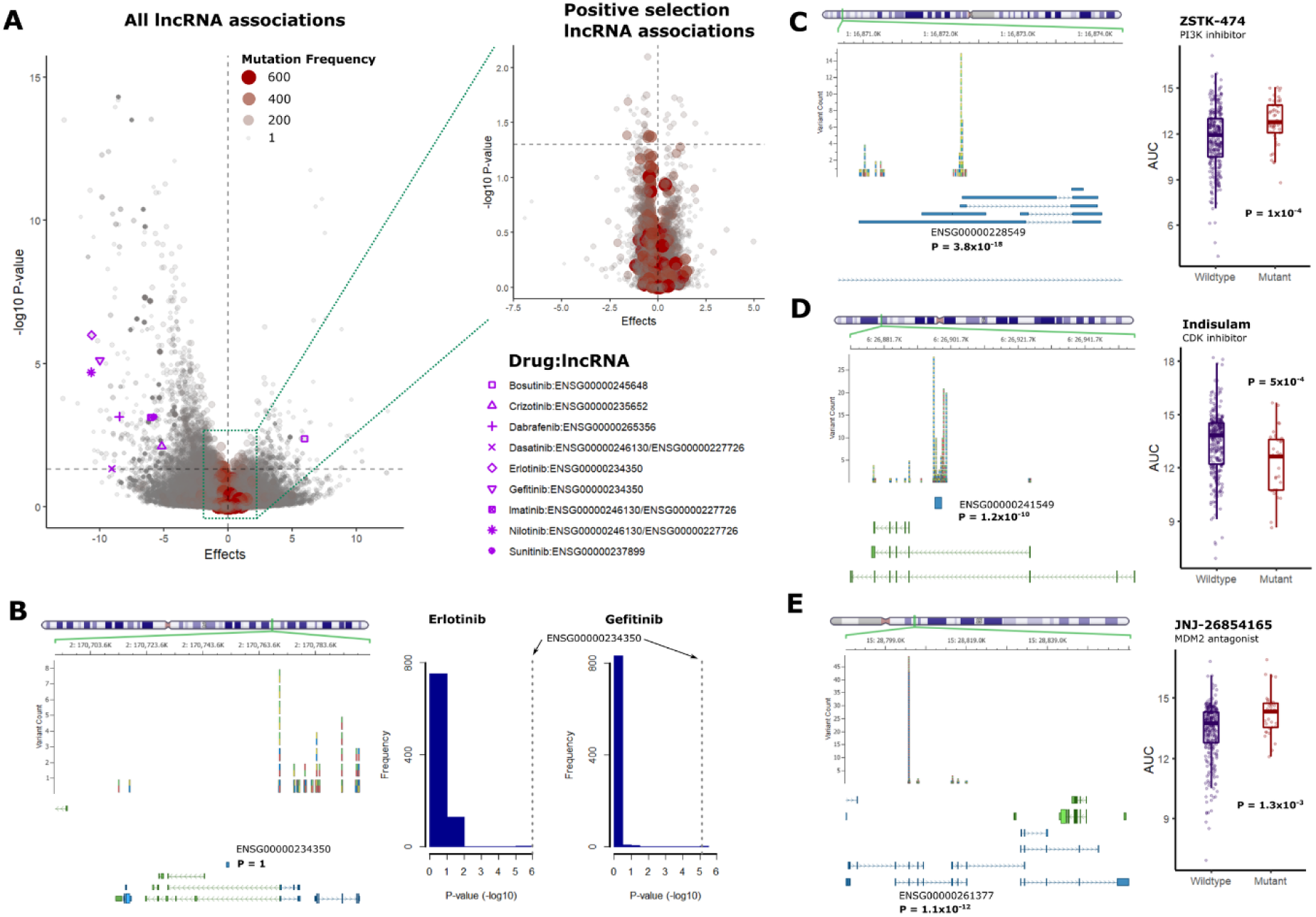
lncRNA-specific somatic variations and association with drug response. **A.** Volcano plot of drug-lncRNA associations based on somatic mutations in lncRNA genes, with the size of points scaled according to the frequency of mutations across all cell lines. Nominal P-value thresholds are indicated by the horizontal dashed grey lines. The inset volcano plot shows lncRNAs satisfying the statistical threshold for positive selection in the COSMIC cell lines. Significant associations above the nominal threshold for drugs with clinically actionable PCG biomarkers are indicated with blue markers. **B.** The left panel shows chromosomal locus of the candidate lncRNA and frequency of non-coding variants identified in the COSMIC cancer cell lines. The exon structure of the proximal PCGs (green) and lncRNAs (blue) are displayed below the variants. The right panel shows the P-value distribution of the lncRNAs associated with erlotinib and gefitinib sensitivity. **C-E.** Examples of lncRNAs with mutation frequencies above the statistical threshold for positive selection in the COSMIC cell lines, with the left panels showing frequency of non-coding variants and exon structure of proximal genes, and right panels showing the comparison of drug AUC in wild-type or mutant cell lines.

We next determined associations for the lncRNAs with somatic mutation frequencies above the statistical threshold for positive selection (Figure 5A inset). As an example, the intergenic lncRNA *BX284668.2* (ENSG00000228549) (P = 3.8×10^−18^) was associated with sensitivity to the PI3K inhibitor ZSTK-474 (P = 1×10^−4^) (Figure 5C). Similarly, mutations in the glucuronidase beta pseudogene 2 (ENSG00000241549) (P = 1.8×10^−10^) were linked with resistance to the CDK inhibitor indisulam (P = 5×10^−4^)(Figure 5D). Additionally, mutations in the programmed cell death 6 interacting protein pseudogene 2 (ENSG00000261377) were determined as positively selected in the COSMIC cell lines (P = 1.1×10^−12^) and predicted sensitivity to the MDM2 antagonist JNJ-26854165 (P = 0.001) (Figure 5E). While the biological function of these lncRNAs is virtually unknown, these examples focusing on lncRNAs undergoing positive selection in cancer cells hint at the existence of PCG-independent associations between somatic alterations in the lncRNA genome and drug sensitivity. At the very least, it is clear that further studies are warranted to characterize the function of somatic lncRNAs variants and study their associations with anticancer drug response.

### Experimental validation of the influence of *EGFR-AS1* and *MIR205HG* expression on erlotinib response in lung cancer cells

We determined the correlation between *EGFR-AS1* and *MIR205HG* expression with erlotinib response in a cohort of 16 lung cancer cell lines (Supplementary Figure 6A). As expected, the cell lines with *EGFR* activating mutations (exon 19 deletions) including HCC 4006, HCC 2935 and HCC 827, were most responsive to erlotinib treatment (Figure 6A). Our experiments yielded similar results as observed in the cancer cell line screens, with higher expression of both lncRNA transcripts associated with increased sensitivity to erlotinib (*EGFR-AS1* PCC = −0.34, *MIR205HG* PCC = −0.26) (Figure 6A).

**Figure 6:**
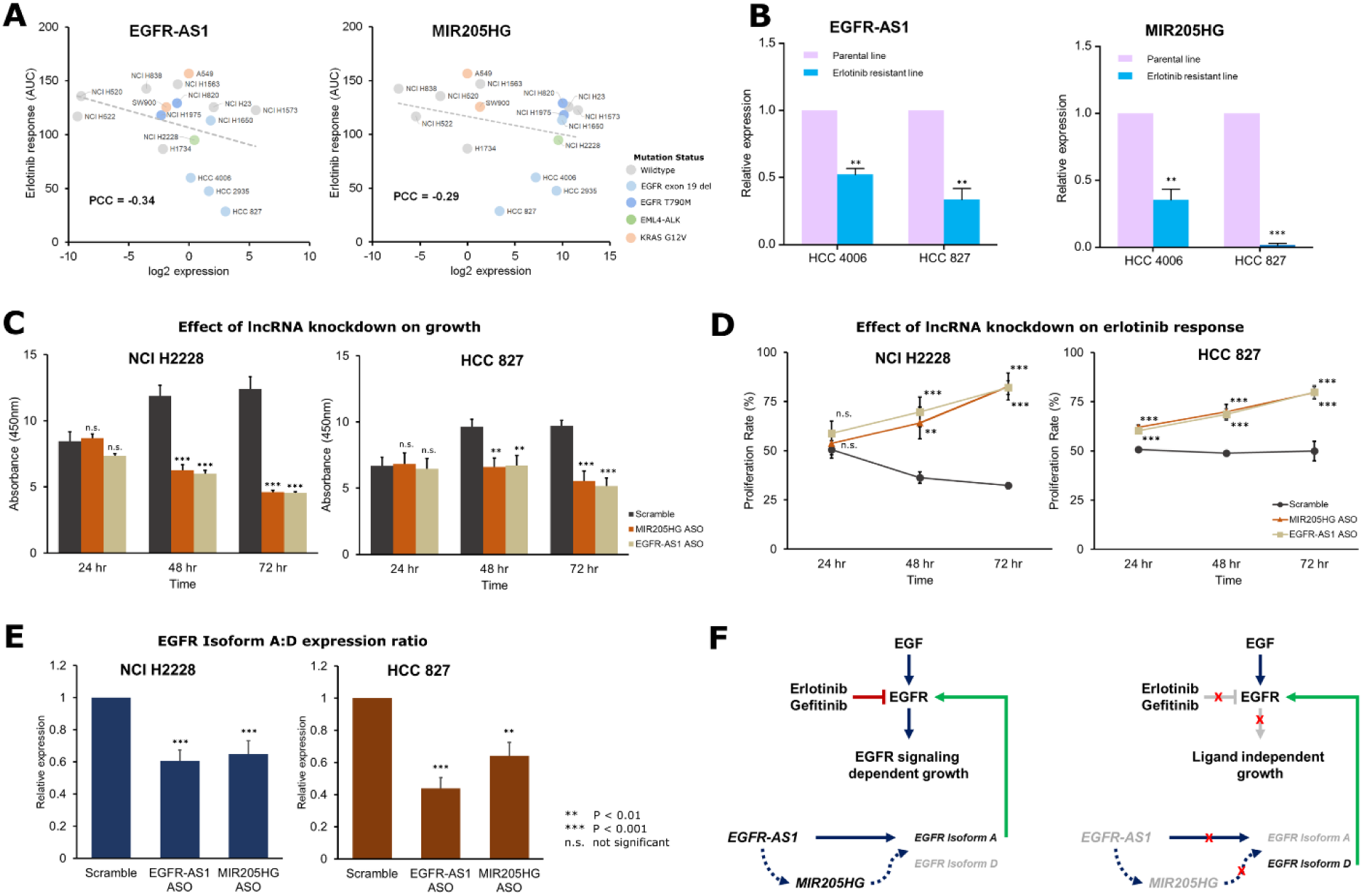
*In vitro* validation of *EGFR-AS1* and *MIR205HG* as determinants of erlotinib response. **A.** Scatter plots showing expression levels of *EGFR-AS1* and *MIR205HG* along with erlotinib response (AUC) for 16 lung cancer cell lines labeled on the plot. Cell lines carrying EGFR-activating mutations (L858R, exon 19 del) are encircled in blue, while lines with resistance mutation (T790M) are encircled in red. The dashed grey line indicates a linear fit. **B.** Comparison of relative *EGFR-AS1* and *MIR205HG* expression levels in two EGFR-responsive cell lines – HCC 4006 and HCC 827. Blue bars indicate expression levels in parental lines, while red bars indicate expression levels in derived, erlotinib-resistant, lines. **C.** Barplots indicating the growth of NCI H2228 and HCC 827 cells upon ASO-mediated k.d. of *EGFR-AS1* or *MIR205HG* over a period of 72 hours post k.d. **D.** Effect of ASO-mediated *EGFR-AS1* or *MIR205HG* k.d. on erlotinib response determined as proliferation rate relative to untreated NCI H2228 and HCC 827 cells measured over a period of 72 hours. **E.** The ratio of relative expression levels of EGFR Isoform A: Isoform D in NCI H2228 and HCC 827 cells with ASO-mediated *EGFR-AS1* or *MIR205HG* k.d. **F.** Schematic of the proposed lncRNA-driven pathway that determines response to erlotinib in the presence (left panel) or absence (right panel) of *EGFR-AS1* and *MIR205HG* mediated regulation of EGFR isoforms.

Next, we measured the expression of the two lncRNAs in two erlotinib-resistant lines generated from erlotinib-sensitive parental lines (HCC 4006 and HCC 827) carrying EGFR-activating mutations (Figure 6B). The expression levels of *EGFR-AS1* were about 50-60% lower in the resistant lines as compared to the parental HCC 4006 and HCC 827 cell lines (both P < 0.01). Similarly, *MIR205HG* expression levels were 60-90% lower in the resistant lines (HCC 4006 P < 0.01; HCC 827 P < 0.001). Based on these results, we hypothesized that these two lncRNAs have a functional influence on erlotinib response and not merely serve as predictive biomarkers. To confirm their functional impact, we performed anti-sense oligonucleotide (ASO) mediated knockdown (k.d.) of *EGFR-AS1* and *MIR205HG* in two cell lines with different levels of erlotinib sensitivity – HCC 827 (strong response) and NCI H2228 (weak response) (Supplementary Figure 6B and 6C). The k.d. of both transcripts resulted in a reduction in the growth of the two cell lines over time, with significantly slower growth observed at 48 and 72 hours (Figure 6C). However, in the presence of erlotinib, the rates of proliferation were elevated in both the *EGFR-AS1* and *MIR205HG* k.d. cell lines as compared to control (Figure 6D). These results corroborate the outcome of our analyses and hint at a possible functional impact of the two lncRNAs on erlotinib response.

To further investigate its functional role, we investigated the relative expression levels of two *EGFR* isoforms that were recently described as regulatory targets of *EGFR-AS1* (Tan et al. 2017). The relative ratio of these isoforms could affect the ligand-dependent activation of the EGFR signaling pathway (Reiter et al. 2001). The product of *EGFR* transcript 1 (ENST00000275493.6 or NM_005228.5) translates into the full-length Isoform A of the protein while transcript variant 4 (ENST00000344576.6 or NM_201284.1) translates into the truncated Isoform D of the protein. Only the extracellular domain is present in the shorter isoform and lacks the tyrosine kinase domain. Thus, abundant expression Isoform D may act as an antagonist of ligand-dependent EGFR action.

In the *EGFR-AS1* and *MIR205HG* k.d. cell lines, the expression levels of the consensus EGFR sequence were not significantly altered (Supplementary Figure 6D). While the Isoform A levels were lower, the reduction was not statistically significant in either lncRNA k.d. in each cell line (Supplementary Figure 6E). The expression levels of Isoform D were elevated in both EGFR-AS1 and MIR205HG k.d. cell lines, but were significant only in HCC 827 cells (Supplementary Figure 6F). However, upon considering the ratios of Isoform A to Isoform D, we found a significant reduction in the relative abundance of the two isoforms in both NCI H2228 (*EGFR-AS1* k.d. P = 1×10^−4^; *MIR205HG* k.d. P = 1.6×10^−3^) and HCC 827 (*EGFR-AS1* k.d. P = 1.6×10^−6^; *MIR205HG* k.d. P = 3.8×10^−7^) cells (Figure 6E). These results indicate a reduction in ligand-dependent growth of the cells and, consequently, reduced impact of erlotinib on the cell lines with lncRNA k.d. (Figure 6F). It is interesting to note that the *MIR205HG* k.d. also resulted in a similar phenotypic impact as *EGFR-AS1* k.d. along with reduced Isoform A:D ratio. Moreover, we observed a significant reduction in *MIR205HG* expression in the *EGFR-AS1* k.d. cells, hinting at a possible mediatory role of *MIR205HG* that calls for further investigation.

## Discussion

Although representing over 80% of the human transcriptome, the pharmacogenomic relevance of lncRNAs in drug response is largely unknown. This gap in knowledge motivated us to comprehensively study the lncRNA transcriptome and genome to determine their relevance in cancer pharmacogenomics. Recent large-scale drug screening efforts (Barretina et al., 2012; Basu et al., 2013; Garnett et al., 2012; Iorio et al., 2016b; Seashore-Ludlow et al., 2015) generated invaluable response data for hundreds of drugs measured across over a thousand cell lines, along with exome sequencing and PCG expression data. We leveraged these datasets to perform an in-depth analysis of the relationship among the lncRNA transcriptome, genome and response to drugs. Previously, expression and somatic alterations of PCGs were successfully utilized to predict drug response using various machine-learning approaches (Costello et al., 2014). From the early drug screens and subsequent prediction efforts, it is clear that the tissue lineage of cancer cell lines has a strong confounding effect on the drug response prediction models (Barretina et al. 2012). Considering the tissue-specific expression patterns of lncRNAs (Jiang et al. 2016), we emphasized the inclusion of tissue type of the cell lines as a covariate in all of our analyses. Moreover, we addressed the unique challenge of identifying drug response-related lncRNAs that are independent of the effects of proximal *cis*-PCGs and well-established PCG biomarkers. Together with confounding effects of tissue-type, adjusting for effects of *cis*-PCGs or known biomarkers is of critical importance in determining potential lncRNA biomarkers. These factors were not taken into account in previous attempts at predicting drug response using lncRNAs (Wang et al. 2018).

In this study, we first analyzed over 3.3 million drug-lncRNA expression associations in the CTRP screen and about 2.3 million associations in the GDSC screen. We determined a large proportion of significant drug-lncRNA associations overlapped in the two independent screens. Furthermore, we hypothesized and validated that the lncRNA transcriptome could effectively predict drug response with equal efficacy as PCGs in both screens.

The physical proximity of a portion of lncRNAs to PCGs and redundant expression patterns raise concerns whether lncRNAs actually provide any additional information. We addressed this important concern by carefully accounting for the possible effects of neighboring PCGs. The choice of ±500Kb boundary that defined *cis*-PCGs in our study was based on the definition of *cis*-eQTLs from the genotype tissue expression project (Lonsdale et al. 2013). Based on this conditional analysis, we determined about half of the initially observed drug-lncRNA associations were redundant with *cis*-PCG expression. However, for most drugs, the genomic loci for the top lncRNA biomarker do not overlap with top the PCGs, including for the drugs with established clinical biomarkers.

Among the known oncogenic and tumor suppressor lncRNAs, we found intriguing associations between *GAS5* and *ZEB2AS1* expression with >50 drugs in both GDSC and CTRP. The *GAS5* (growth arrest-specific 5) lncRNA acts as a tumor suppressor with various proposed mechanisms of action, including, cell cycle control (Liu et al. 2015; Mazar et al. 2016; Luo et al. 2017), proliferation (Kino et al. 2010; Li et al. 2017b) and regulation of epithelial to mesenchymal transition (EMT) program (Zhuang et al. 2015). Considering the multi-faceted mechanisms by which *GAS5* functions as a tumor suppressor, it is plausible that its expression may be predictive of response to multiple drugs. In contrast, from what we know so far about *ZEB2AS1*, it appears this transcript upregulates *ZEB2* expression to induce EMT program in cancer cells (Beltran et al. 2008; Zhuang et al. 2015). One possible mechanism could be the expression of this transcript is indicative of aggressive cancer cells that generally show a better response to drugs *in vitro*. Nevertheless, both lncRNAs are candidates for experimental validation as multi-drug response prediction biomarkers.

In addition to the lncRNA transcriptome, we study the associations between somatic alterations in the lncRNA region and drug response. Currently, there are no gold standards to distinguish driver vs. passenger non-coding somatic variants. Unlike PCGs, non-coding variants cannot be classified as synonymous or non-synonymous, which complicates identification of relevant variants. We attempted to identify lncRNA variants that did not overlap with any PCGs and were positively selected in the cell lines based on the total background non-coding mutation frequency and lncRNA mutation frequency across all cell lines. While preliminary, the emergence of significant somatic lncRNA associations with drug-response warrants future studies focusing on elucidating the biological role of such variants.

In clinical practice, a handful of somatic variants are routinely profiled to guide treatment decisions in cancer patients (Relling and Evans 2015). Among these, somatic mutations that activate EGFR activity and their impact on anti-EGFR therapeutics like erlotinib and gefitinib are some of the first and extensively studied clinically actionable biomarkers (Jimeno and Hidalgo 2006). An important consideration in identifying novel lncRNA biomarkers is accounting for such well-known PCG somatic alterations to prevent redundancy. In this direction, we identified and characterized the impact of two lncRNAs, *EGFR-AS1*, and *MIR205HG*, which are strong predictors of response to erlotinib and gefitinib independent of *EGFR* somatic mutation status. A recent study proposed the theory that *EGFR-AS1* stabilizes the EGFR signaling pathway by influencing the transcript ratio of the full-length EGFR Isoform A to the truncated Isoform D (Tan et al. 2017). Our results lend support to this idea, demonstrating the effect of *EGFR-AS1* k.d. on the ratio of the two isoforms. Moreover, we reported a shift in both the growth pattern and erlotinib sensitivity of cell lines upon *EGFR-AS1* k.d. confirming the functional impact of this lncRNA on addiction to ligand-dependent EGFR signaling. The *MIR205HG* lncRNA undergoes post-transcriptional processing leading to the synthesis of miRNA-205. Several studies have focused on the functions of this miRNA, however, its mechanism of action remains conflicting, with both oncogenic and tumor suppressor activities proposed (Greene et al., 2014; Niu et al., 2015). Similar to *EGFR-AS1*, the k.d. of *MIR205HG* also resulted in a change in *EGFR* isoform ratio and phenotypic outcome. Additionally, *MIR205HG* expression levels were affected by *EGFR-AS1* k.d., suggesting this gene could be an intermediator in EGFR isoform regulation. While miRNA-205 does not bind and regulate *EGFR* expression directly, it could modulate secondary transcriptional repressors that regulate the expression of the transcripts (De Cola et al. 2015) (Figure 6F). Besides an intriguing functional link with EGFR signaling, the expression of these two lncRNAs could predict response in patients who do not carry known activating *EGFR* mutations and thus are strong candidates for clinical evaluation.

## Conclusions

We are still in the early stages of understanding the biology of lncRNAs, though sufficient evidence points towards the direction of altered lncRNA’s involvement in human cancers. By comprehensively analyzing the associations between lncRNA transcriptome, genome, and drug response, we have shown that lncRNAs are indeed biomarkers of drug response. Furthermore, we have provided compelling evidence that these associations are not just dependent on the correlative structure with PCGs. Future studies focusing on the mechanism of lncRNA action would be invaluable in improving our understanding of cancer progression and drug response.

## Materials and Methods

### Gene expression datasets and lncRNA imputation

The original GDSC, CCLE and CTRP studies utilized microarrays to profile the transcriptome of cancer cell lines and identify associations between expression and drug response. In order to facilitate the analysis of lncRNA transcriptome with drug response, we developed a lncRNA expression imputation framework (LEXI) that enables accurate imputation of lncRNAs in uncharacterized samples only using PCG profiles (Nath et al. 2019). Briefly, our approach harnesses redundancy in PCG and lncRNA expression profiles to generate imputation models using machine learning. These models can then be implemented to generate lncRNA transcriptome of samples with only PCG data. We downloaded GDSC microarray gene expression data from https://www.cancerrxgene.org/, also available from ArrayExpress (E-MTAB-3610), normalized using the RMA method. The PCG profiles from this dataset served as the training set for LEXI. We also obtained gCSI RNAseq gene expression data (RPKM) for 675 cell lines available at http://research-pub.gene.com/KlijnEtAl2014/ and ArrayExpress (E-MTAB-2706). We annotate genes from the gCSI RNAseq dataset based on GENCODE (release 28) biotypes to classify genes as either PCGs or lncRNAs. PCGs were defined as genes with biotype “protein coding” and lncRNAs were defined as genes with transcripts products of length >200nt and including the biotypes “3prime_overlapping_ncrna”, “antisense_RNA”, “bidirectional_promoter_lncRNA”, “lincRNA”, “processed_transcript”, “retained_intron”, “sense_intronic”, or “sense_overlapping”. The PCG and lncRNA expression profiles from gCSI served a training dataset for imputing lncRNA transcriptome of GDSC cell lines.

### Predicting drug response using PCG or lncRNA expression

We obtained drug response data from GDSC (https://www.cancerrxgene.org/) and CTRP (https://portals.broadinstitute.org/ctrp/) portals for 265 and 545 anticancer agents respectively. Additional drug and cell line annotation regarding targets/pathways, disease and tissue-type were retrieved from the respective portals. In this study, we utilized the area under the curve (AUC) parameter from the dose-response data to construct a response vector Y for each drug tested in n cell lines. Next, we constructed prediction feature matrices of gene expression X measured in n cell lines. The feature matrix was constructed separately for PCGs and lncRNAs. For the 963 GDSC cell lines, we used the measured microarray PCG expression data and lncRNA transcriptome imputed using gCSI as training data. We also used the subset of cell lines that overlapped between GDSC and gCSI sequencing data to compare the efficiency of predicting response using the measured lncRNA transcriptome. For the CTRP cell lines, we used measured PCG and lncRNA expression levels (RPKM) from RNAseq data of 1019 cell lines from CCLE (https://portals.broadinstitute.org/ccle/). For each dataset, we used the definition of PCG and lncRNA as defined above in the imputation section based on GENCODE annotation and transcript product length. The training matrices were processed to remove all features with variance = 0, and standardized such that each gene expression vector had a mean = 0 and standard deviation = 1. The model for the prediction analysis can be expressed in the form of the following linear equation:

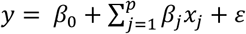

Where,

*y* = *AUC vector of the drug measured in* “*n*” *samples*, *β*_0_ = *intercept*, *β_j_* = *coefficient associated with gene_j_*,

*x_j_* = expression vector of gene_j_, *and* ε = *model error*

Since the number of features is much greater than the number of samples (p >> n), we used ridge regression for AUC prediction by cross-validation as this method was found to consistently outperform other methods with high computational efficiency in predicting drug sensitivity (Geeleher et al. 2014; Azuaje 2016). We fit the ridge model using PCG or lncRNA expression matrices as features by solving the following objective function using the R-package glmnet (Friedman et al. 2010):

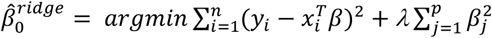

Where,

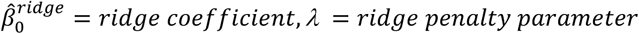

Here, the penalty parameter is determined by performing 10-fold cross-validation on the training dataset. Following the determination of optimal lambda, we use the cross-validation to determine prediction accuracy by calculating the average Spearman’s correlation coefficient between predicted and actual AUC, and root means squared error (RMSE) of the model. The significance of prediction is determined based on a null distribution of prediction accuracies generated from the feature set of 5×10^6^ randomized gene expression and 5×10^3^ AUC values for 1000 samples, and correlations with P < 0.05 are considered significant.

### Determining significant lncRNAs associated with drug response

Previous pharmacogenomic analyses in cancer cell line screens have revealed that the drug response is strongly linked with cell line lineages. In addition, lncRNAs are understood to be expressed in a highly tissue-specific manner. To address both these issues while identifying novel lncRNA transcripts that are associated with drug sensitivity or resistance, we use similar linear model used for drug prediction as above but with the addition of a covariate to adjust for tissue type:

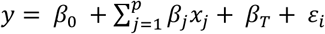

Where,

*β*_T_ = *Coefficient for tissue type*

In addition, we identified lncRNAs biomarkers that are independent predictors of drug response from known PCG biomarkers. We utilized the cancer function events (CFE) information from (Iorio et al. 2016a), which included high-confidence cancer genes from the intOGen pipeline and (Wong et al. 2013), recurring copy number alterations (van Dyk et al. 2013) and CpG islands. We used the binary event matrix of CFEs across all cell lines, and individual events that are associated with response to a specific drug were included in the model:

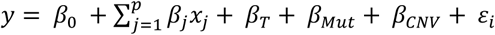

Where,

*β_Mut_* = *Coefficient for mutation CFE*

*β_CNV_* = *Coefficient for copy number vairation CFE*

For example, for the EGFR-targeting drugs erlotinib, gefitinib and pelitinib, we used mutations and amplification of the EGFR gene as CFEs. We solve the above two equations using multiple linear regression to obtain the coefficients and p-value associated with each drug-gene pair. Since the number of multiple comparisons is in order of 10^6^, we used a Bonferroni adjusted threshold to determine statistically significant lncRNA-drug pairs.

### Determining the impact of proximal *cis*-PCGs on drug-lncRNAs associations

We defined *cis*-PCGs as genes located within a 1 Mb boundary of the lncRNA transcript, i.e. all PCGs that start or end within 500kb upstream of the lncRNA transcription start site to 500kb downstream of the lncRNA transcript end. This definition of *cis*-PCGs is derived from the 1 Mb boundary used to determine *cis*-eQTLs in the GTEx project (Lonsdale et al. 2013). The genomic and transcript coordinates of lncRNAs and PCGs were based on the human genome reference assembly GRCh38.p12. We analyzed the impact of *cis*-PCGs on significant drug-lncRNA associations by including the expression levels of all *cis*-PCGs for the lncRNA in the model:

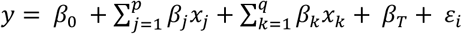

Where,

*β*_k_ = *Coefficient associated with the expression cisPCG x_k_*

The p-values (−log_10_) of the significance of association for each individual variable after adjusting for the covariates were deemed significant if P < 0.05 and mapped based on chromosomal coordinates using the R/Bioconductor package ‘karyoploteR’ (Gel et al. 2017).

### LncRNA-specific somatic mutations

A number of statistical tools have been developed to identify driver somatic mutations in cancer cells. The general objective of these methods is identifying non-random mutations with functional impact on proteins of interest, usually relying on a statistical model that estimates background mutation rates based on distribution of synonymous and non-synonymous alterations (Dees et al. 2012; Gonzalez-Perez and Lopez-Bigas 2012; Lawrence et al. 2013; Reimand et al. 2013). However, these methods, by definition, cannot infer somatic mutations in the non-coding genome. Recent efforts have been directed toward identifying somatic drivers that are specifically located in the lncRNA genome (Li et al. 2015; Lanzos et al. 2017). To identify lncRNA genes with positively selected somatic mutations in the cancer cell lines, we adopt a two-step approach that first selects lncRNA genes with variation frequency higher than the non-coding genome background across all available cell lines, and second, scores lncRNA genes based on mutation frequency per cell line. While highly stringent, this approach can be expected to yield high-confidence somatic variants in the lncRNA region.

The somatic mutations used in our study were called from whole genome sequencing data of 1015 COSMIC cell lines (https://cancer.sanger.ac.uk/cell_lines/download) using the Caveman and Pindel algorithms (Iorio et al. 2016a). The variants were additionally filtered based on read-depth (≥15), mutant allele frequency (≥15%) and absence from reference normal samples, 1000 genomes, and dbSNP. We annotated the remaining variants based on the GRCh38.p12 human reference genome and GENCODE 28 gene set using the Variant Effect Predictor tool (McLaren et al. 2016). We removed all variants that mapped on regions annotated as protein-coding genes or within genes that encoded transcripts less than <200nt in length. We further removed ambiguous variant locus overlapping protein-coding and lncRNA genes.

We first identify lncRNA genes with a mutation frequency by solving the following equation for the Kolmogorov-Smirnov test (α = 0.05) for each *n* lncRNA, considering non-coding mutation for all *m* genes as background:

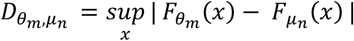

Where,

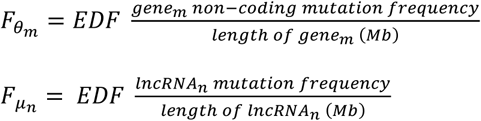

(EDF, empirical distribution function)

Next, we use a one-tailed hypergeometric distribution to test the probability whether a given lncRNA *n* is mutated at a greater frequency than the background frequency of all lncRNAs, at a significance threshold of p < 0.05, using the equation:

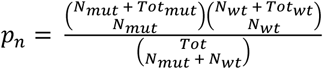

Where,

*N_mut_* = *Number of cell lines with lncRNA_n_ mutations*,

*N_wt_* = *Number of cell lines without lncRNA_n_ mutations*,

*Tot_mut_* = *Total number of cell lines with any lncRNA mutations*,

*Tot_wt_* = *Total number of cell lines without any lncRNA mutations*,

*and, Tot* = *Tot_mut_* + *Tot_wt_*

The lncRNAs with somatic mutation frequencies above the background level were retained for further analysis using multiple linear regression analysis against drug response yielding coefficients and a p-value of association.

### TCGA lung adenocarcinoma (LUAD) analysis

RNAseq (RSEM) expression levels of *EGFR-AS1* and *MIR205HG*, and EGFR point mutation, copy number variation data, and survival data for 877 lung adenocarcinoma (LUAD) samples were retrieved using UCSC Xena platform for cancer genomics (Goldman et al. 2018). Imputed erlotinib drug sensitivities for TCGA LUAD were obtained from (Geeleher et al. 2017). Correlation, multiple regression, and ANOVA analyses were performed with or without including *EGFR* somatic mutations as covariates using either continuous expression data or samples clusters (K-means) based on expression levels of the lncRNAs. Kaplan-Meir (survival) curves were analyzed by the log-rank (Mantel-Cox) test using the ‘survival’ R package.

### Mammalian cell culture and reagents

The human lung cancer cell lines A549, NCI H1563, NCI H1573, NCI H1650, NCI H1734, NCI H1975, NCI H2228, NCI H23, NCI H520, NCI H522, NCI H820, NCI H838, HCC 2935, HCC 4006, HCC 827, SW900 were purchased from American Type Culture Collect (ATCC). A549 cells were cultured in DMEM-HG, and the other cell lines were cultured in RPMI 1640 medium (Thermo Fisher Scientific) supplemented with 10% fetal bovine serum (FBS) (Gemini Bio-Products) and 1% penicillin and streptomycin (Life Technologies), maintained at 37°C in a humidified incubator with 5% CO2 atmosphere. Cell lines were observed microscopically to confirm morphology, and population-doubling times were determined by viable cell counting using trypan blue on the TC20™ Automated Cell Counter (Bio-Rad). Aliquots of low-passage cells were cryopreserved within 2 weeks of receipt. Cells were cultured for no longer than 10 total passages. All cell lines were periodically monitored for mycoplasma using the Universal Mycoplasma Detection Kit following the manufacturer’s protocol (ATCC). Culture health and identity were monitored by microscopy and by comparing population doubling times to baseline values recorded at the time of receipt.

### Generating erlotinib-resistant lung cancer cell lines

Erlotinib resistant HCC 827 (harboring EGFR exon 19 deletions) and HCC 4006 (harboring EGFR exon 19 deletions) cell lines were generated by chronic exposure of the cells to a stepwise increasing dose of erlotinib to 3μM and 9μM respectively for up to 6 months.

### LncRNA knockdown using 2’-deoxy-2’-fluoro-arabinoguanosine (FANA) antisense oligonucleotides (ASOs)

Knockdown of the *EGFRAS1* and *MIR205HG* lncRNA transcripts was performed FANA ASOs provided by AUM BioTech (AUM BioTech, LLC, Philadelphia, PA). The target sequences for FANA-ASO-*EGFR-AS1* and FANA-ASO-*MIR205HG* were 5’- TATACATTTCATCCCATTGAC-3’ and 5’-AAGATTGAGCCACTGCACTCC-3’, respectively. FANA-ASO scramble served as the control. Knockdown efficiency was experimentally determined by treating HCC827 and NCI H2228 cells with different working concentrations (1µM, 5µM, and 10µM) of FANA ASOs following the manufacturer’s protocol. Cells from both lines were seeded at 5×10^5^ cells across three 6-well culture plates in RPMI 1640 with 10% FBS and maintained at 37°C in a humidified incubator with 5% CO2 atmosphere for 24hrs to allow cells to adhere. After 24hrs, the media was aspirated and replaced with fresh growth media containing the appropriate FANA-ASO at the desired working concentration. Cells were harvested at 24hr, 48hr, and 72hr after the addition of FANA-ASOs to determine knockdown efficiency.

### Quantitative real-time PCR

For the experiments involving FANA-ASO mediated knockdown of *EGFR-AS1* and *MIR205HG* expression, total RNA was extracted from cultured cells using the Quick-RNA MiniPrep Plus Kit (Zymo Research, Irvine, CA) and quantified on the NanoDrop ND-8000 8-channel spectrophotometer (Thermo Fisher Scientific, Waltham, MA). A total of 2µg of RNA was used to synthesize cDNA utilizing the High Capacity cDNA Reverse Transcription Kit (Thermo Fisher Scientific, Waltham, MA). The Sso-Advanced Universal SYBR Green SuperMix (Bio-Rad, Hercules, CA) was used to conduct real-time PCR analyses following the manufacturer’s protocol. The PCR primers used to amplify target lncRNAs and housekeeping genes are as follows: *EGFR-AS1* Forward 5’- GACCACACTGAGCACTCAATAA -3’, *EGFR-AS1* Reverse 5’- CATGCAGCACACACACATTC -3’, *MIR205HG* Forward 5’-GCTCACCCTTGACTTGGAAA -3’, *MIR205HG* Reverse 5’-GGAATTGAAGGAGAGGGAGTAAAG -3’, *EGFR* Forward 5’-GGTGACTCCTTCACACATACTC -3’, *EGFR* Reverse 5’- CCTGCCGCGTATGATTTCTA - 3’, *EGFR* Isoform-A Forward 5’- ACTCTGAGTGCATACAGTGC -3’, *EGFR* Isoform-A Reverse 5’- TCGTTGGACAGCCTTCAAGAC -3’*, EGFR* Isoform-D Forward 5’-ACTCTGAGTGCATACAGTGC -3’*, EGFR* Isoform-D Reverse 5’-TGAAGGCATGAGGCTCAGTG -3’, *GAPDH* Forward 5’- GAACATCATCCCTGCCTCTAC-3’, *GAPDH* Reverse 5’- CCTGCTTCACCACCTTCTT -3’, *ACTB* (*β*-Actin) Forward 5’-GTGGCCGAGGACTTTGATT -3’,and *ACTB* (*β*-Actin) Reverse 5’-TTTAGGATGGCAAGGGAC TTC -3’. Q-real-time PCR and data collection were performed using the 7500 Real-Time PCR System (Applied Biosystems, Foster City, CA). All results were normalized with the expression of *GAPDH* and *β*-Actin as a reference panel. All PCR reactions were performed in triplicate, and each experiment was performed independently three times. Expression results were quantified using the ΔΔCt method relative to the scramble control. For the experiments measuring *EGFR-AS1* and *MIR205HG* expression across 16 lung cancer cell lines and erlotinib-resistant cell lines, total RNA was isolated using Trizol reagent (Life Technologies) according to the manufacturer’s instruction. Complementary DNA (cDNA) was generated with 1μg of total RNA using SuperScript III First-Strand Synthesis System (Thermo Fisher Scientific) according to the manufacturer’s protocol. qPCR and lncRNA expression analysis was performed by the LightCycler® 480 System (Roche). Target lncRNA expression was calculated by the relative quantification method (ΔΔCT) with *RNU44* as the reference control.

### Cell proliferation assays and drug response curves

Lung cancer cell lines were seeded at a density of at 0.5-1×10^4^ cells per well into 96-well plates and incubated for 24 hours at 37°C, followed by exposure to erlotinib at various concentrations for 72 hours. Cell viability was subsequently measured by incubation with WST-1 (Roche Applied Science, Penzberg, Upper Bavaria, Germany) following the manufacturer’s protocol. Following a 2hr incubation at 37°C after WST-1 addition, absorbance at the 450nm wavelength was assessed using the Synergy HTX Multi-Mode Plate Reader (BioTek, Winooski, VT). To assess the impact of ASO-mediated knockdown of the lncRNAs on proliferation and erlotinib response, HCC827 and NCI H2228 cells were cultured in 96-well plates for 24hrs. Next, the cells were treated with media only or FANA-ASO at a working concentration of 5uM, followed by erlotinib treatment approximately 8 hr after FANA-ASO exposure. Cell proliferation was measured at 24hr, 48hr, and 72hr after FANA-ASO +/− erlotinib treatment by the cell proliferation reagent WST-1 as described above. Each experiment was performed in triplicates and measurements were reported as the average of three independent biological replicates.

### Statistical analysis and software

All imputations, predictions and statistical analysis were performed in the R statistical computing environment. For multiple comparisons, family-wise error rates (FWER) were controlled using the Bonferroni method and false discovery rates (FDR) were determined using the Benjamini-Hochberg method where specified. The drug sensitivity curves were plotted and analyzed to obtain the area under the curve (AUC) or IC50 values using GraphPad Prism 6.01. In order to ensure reproducibility and for complete transparency, we have provided links to publicly available datasets. Additionally, we have provided the scripts necessary to perform the analysis available at https://osf.io/wpsdu/ (https://doi.org/10.17605/OSF.IO/WPSDU).

## Declarations

### Author contributions

A.N. conceived the study, collected data and performed the analysis with input from P.G and R.S.H. E.Y.L. and A.L. designed and performed the in vitro experiments with input from A.N., W.S.C. and R.S.H. A.N. wrote the manuscript. All authors provided feedback on manuscript drafts and approved the final version.

### Data and materials availability

All data and materials used in the manuscript are publicly available and listed under methods. The link for the repository with scripts necessary to perform the statistical analyses is also listed in methods.

## Supplementary Materials

**Supplementary Figure 1:**
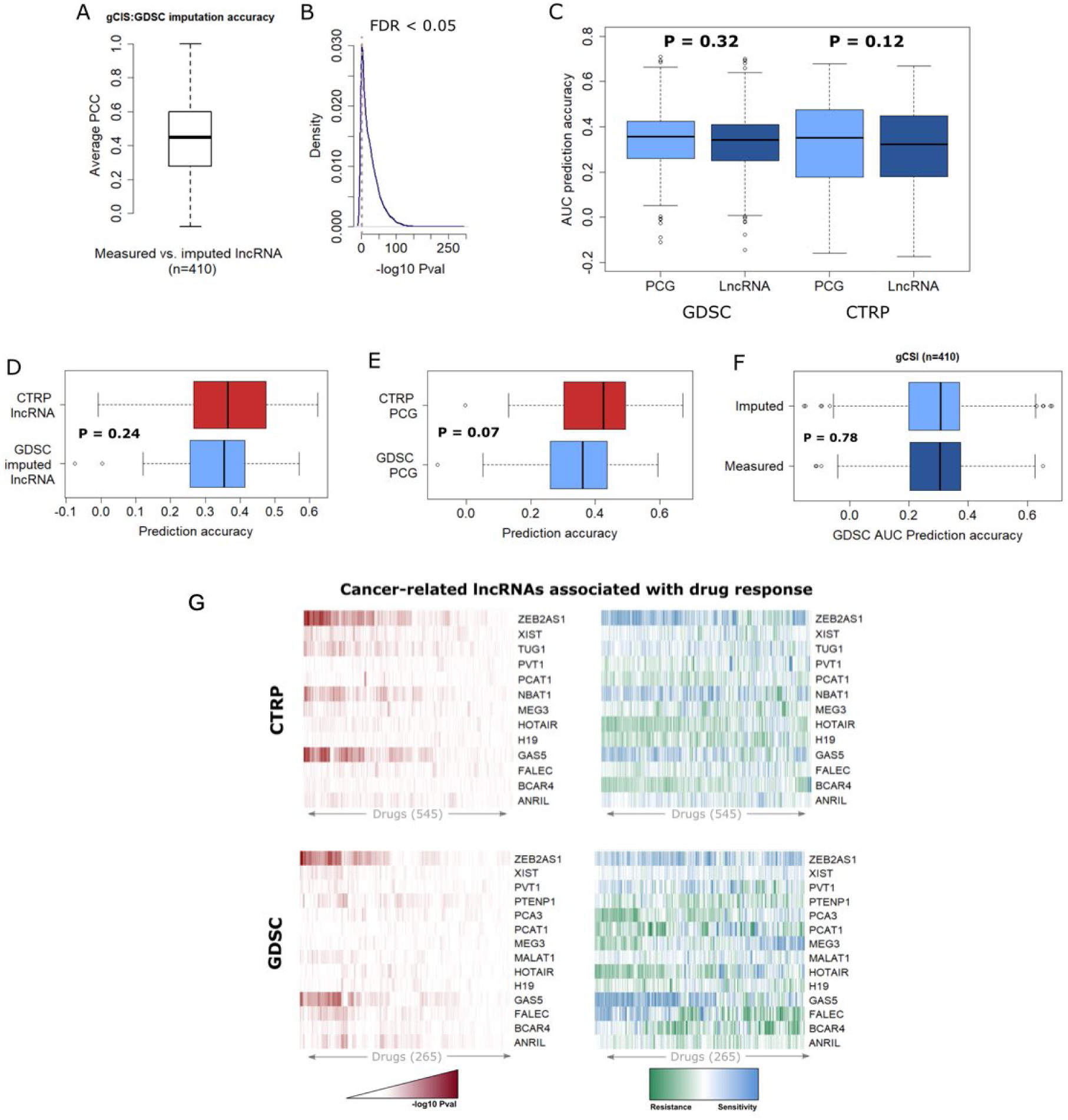
Drug response prediction using lncRNA and PCG transcriptomes. **A.** Boxplot showing the accuracy of imputing lncRNA transcriptome of the GDSC cell lines using the Genentech Cell Screening Iniative (gCSI) RNAseq training dataset. Imputation accuracy is represented as Pearson’s correlation coefficient between measured and imputed lncRNA expression levels in the overlapping cell lines. **B**. Density plot for the significance of the correlation between gCSI measured and imputed lncRNA expression levels in the GDSC dataset. Dashed grey line indicates the threshold of significance (FDR < 0.05). **C.** Boxplots showing the comparison of AUC prediction accuracies between PCG and lncRNA models for the GDSC and CTRP datasets. **D.** Comparison of AUC prediction accuracies of drugs that overlap CTRP and GDSC dataset using measured or imputed lncRNA expression respectively or **E.** PCG expression. **F.** Comparison of GDSC AUC prediction accuracies between imputed lncRNA models from GDSC cell lines and measured lncRNA models from the overlapping set of gCSI cell lines. **G.** Visualization of the significance of association (top panels), direction and magnitude of effects (bottom panels) between CTRP and GDSC drugs with known oncogenic or tumor suppressor lncRNAs. The heatmap shows lncRNAs along Y-axis and different drugs along X-axis. Colors on the top panel are scaled from white to red according to increasing magnitude of –log10 of the p-value, while bottom panel colors scaled from green to blue to indicate whether the lncRNA is correlated with resistance or sensitivity to a drug.

**Supplementary Figure 2:**
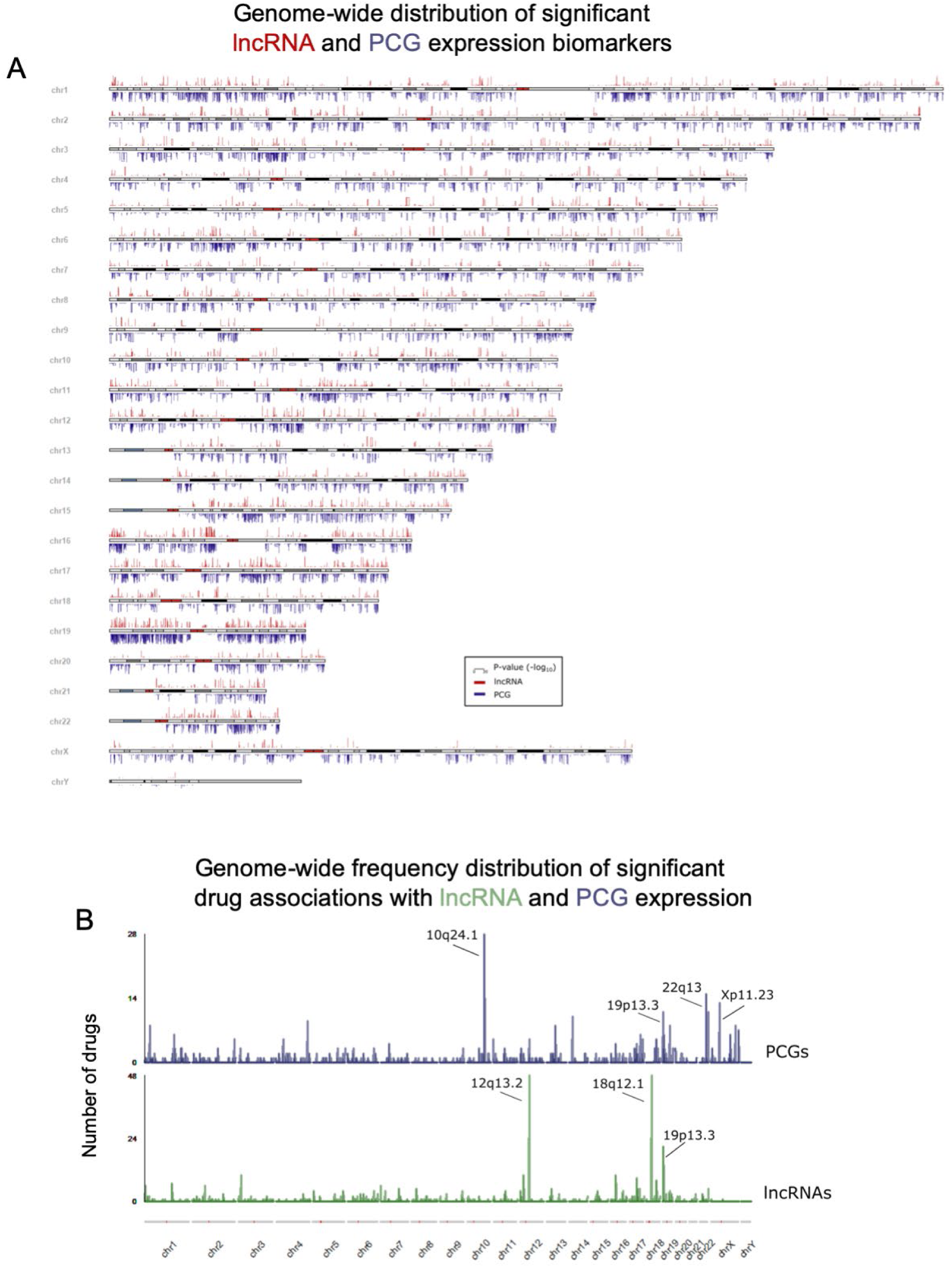
Genomic landscape of drug-lncRNA associations. **A.** Genomic landscape of PCG and lncRNAs associated with CTRP drug response. Each bar indicates the significance of the association between a drug and either PCG (blue bar) or lncRNA (red bar) determined using regression analysis adjusting for tissue type. **B.** Histogram displaying the frequency of significant drugs associated with PCG (blue) and lncRNA (red) loci, grouped by cytobands.

**Supplementary Figure 3:**
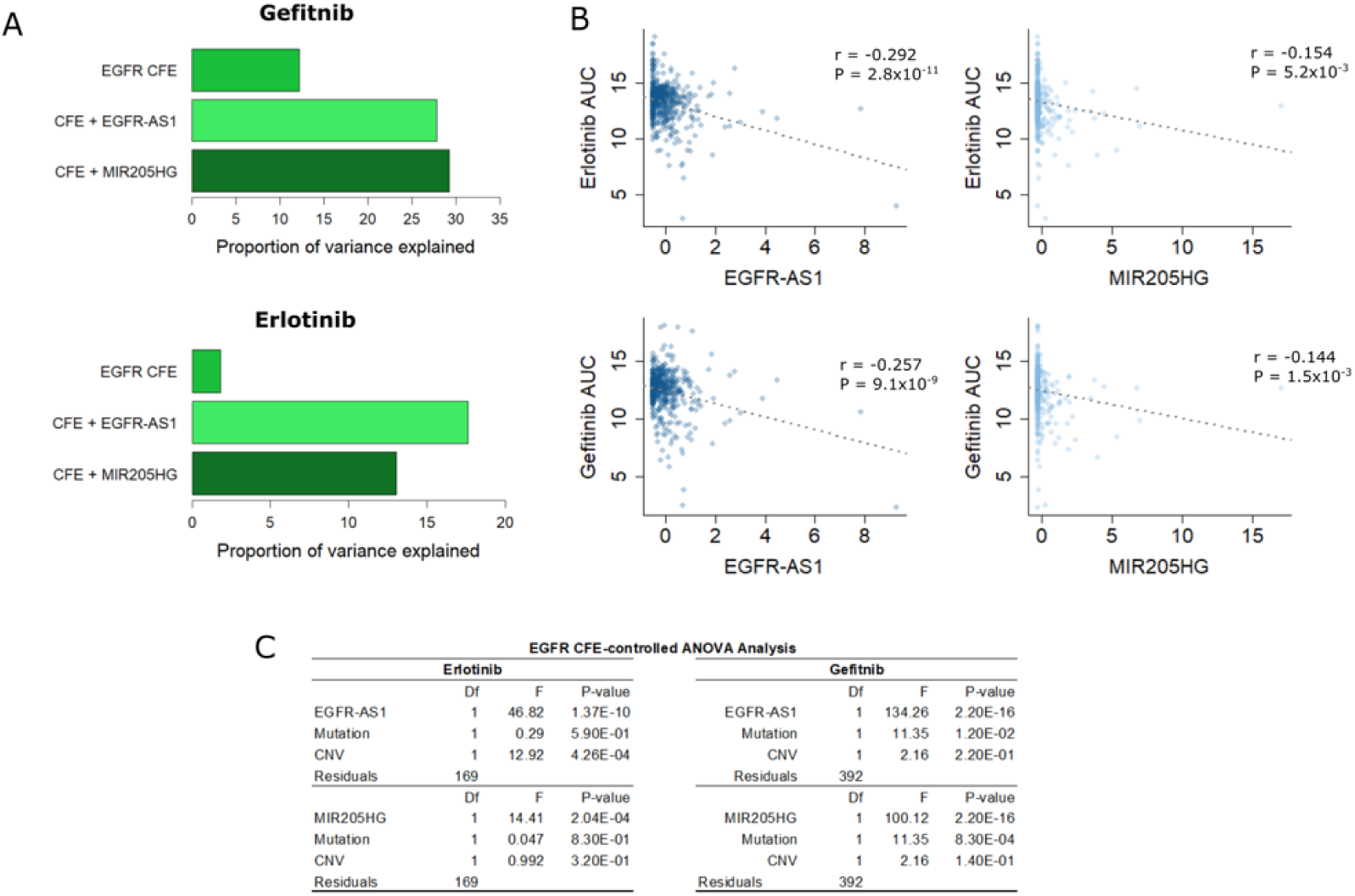
lncRNAs associated with anti-EGFR drugs. **A.** Volcano plot showing lncRNAs associated with pelitinib response in the GDSC dataset adjusting for mutations and copy number variations in the EGFR gene. Each point on the plot represents a lncRNA-drug pair, with blue points indicating lncRNAs common to pelitinib and erlotinib or gefitinib above the nominal significance threshold of FDR < 0.05 (dashed grey line). **B.** Scatter plots showing correlations between measured expression levels of *EGFR-AS1* or *MIR205HG* with erlotinib or gefitinib response (AUC) in the CTRP dataset. **C.** Results of the ANOVA analysis for erlotinib and gefitinib response (AUC) using imputed expression levels of either *EGFR-AS1* or *MIR205HG* while adjusting for EGFR functional events, including mutations and copy number variations.

**Supplementary Figure 4.**
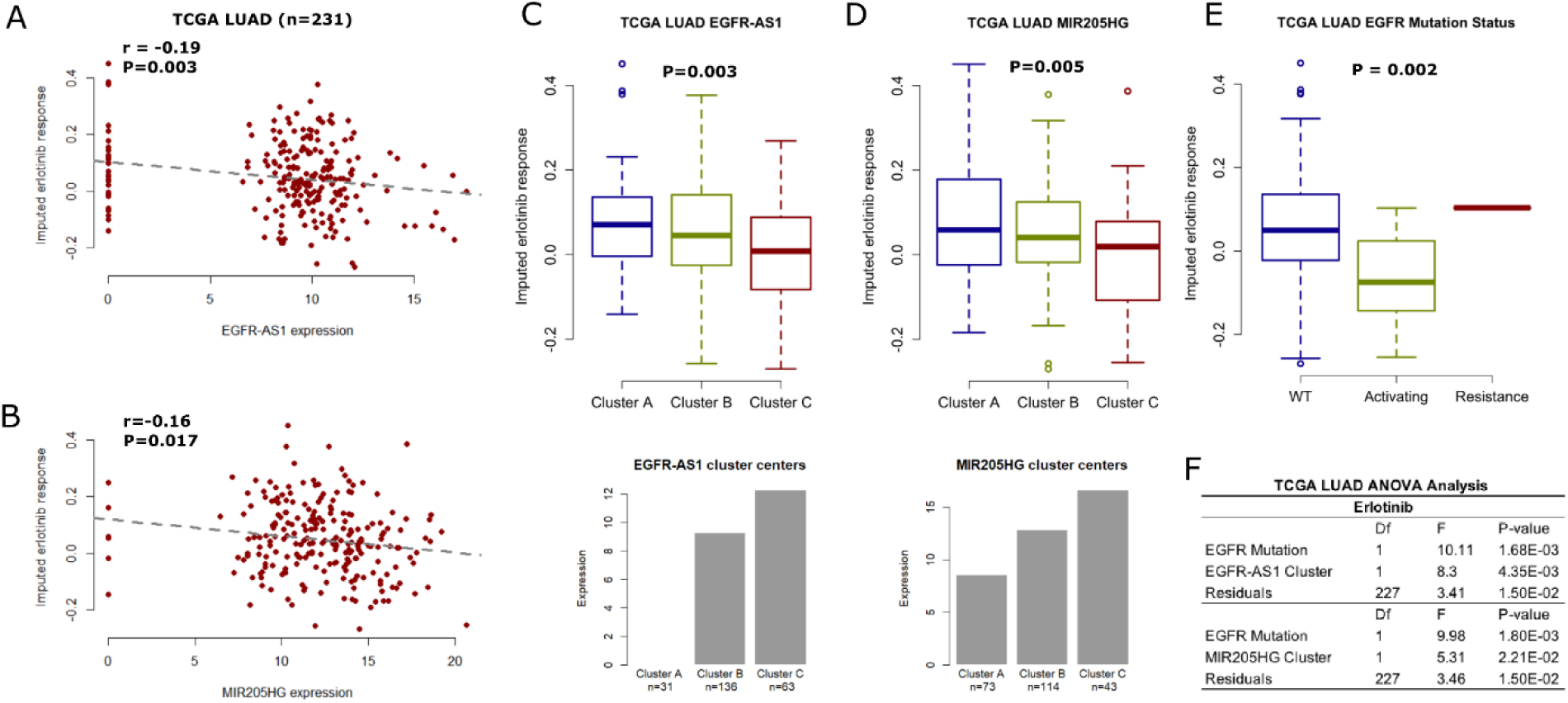
Validation of erlotinib response prediction in TCGA. **A-B.** Scatter plots comparing expression levels of *EGFR-AS1* (A) or *MIR205HG* (B) with imputed erlotinib response in the TCGA LUAD cohort, with higher imputed response scores indicating poor response (similar to high AUC) and lower scores indicating a better response. The dashed grey line indicates a linear fit. **C-E.** Boxplots comparing imputed drug response in the TCGA LUAD patients grouped by *EGFR-AS1* expression clusters (C), *MIR205HG* expression clusters (D) or *EGFR* mutation status (E). The clusters for the two lncRNAs were obtained using k-means clustering, with the centers of the clusters indicated in the bar plots below. **F.** ANOVA analysis of *EGFR* mutation status and lncRNA clusters showing independent associations with imputed erlotinib response

**Supplementary Figure 5:**
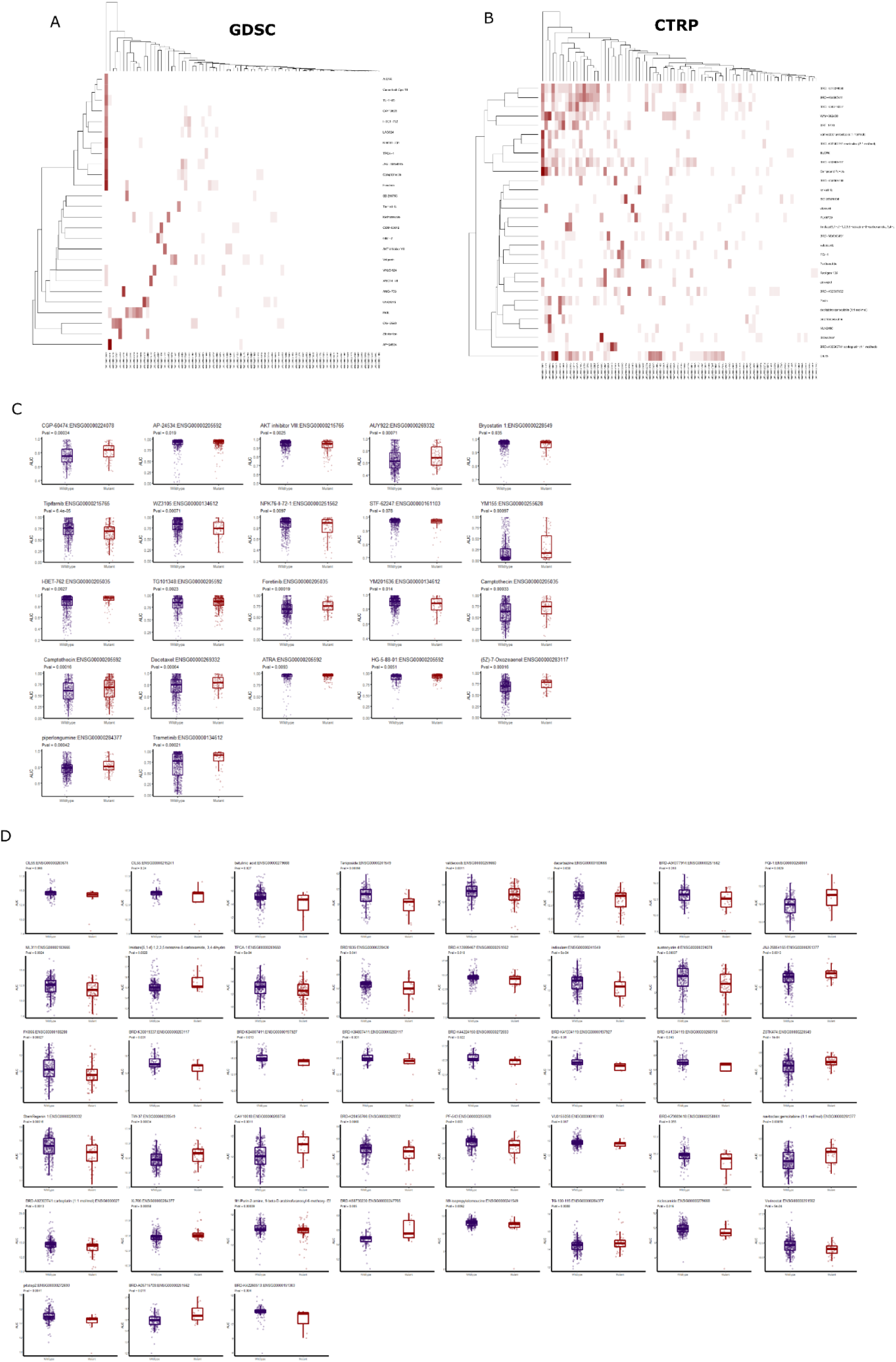
Significant drug-lncRNA somatic mutation associations. **A-B.** Heatmaps significance of the associations (−log10 P-value) between GDSC (A) and CTRP (B) drugs with lncRNAs with mutations above the statistical threshold for positive selection in the COSMIC cell lines. Darker shades of red indicate stronger associations. **C-D.** Boxplots comparing drug AUC in GDSC (C) and CTRP (D) screens based on the mutation status of the lncRNA.

**Supplementary Figure 6:**
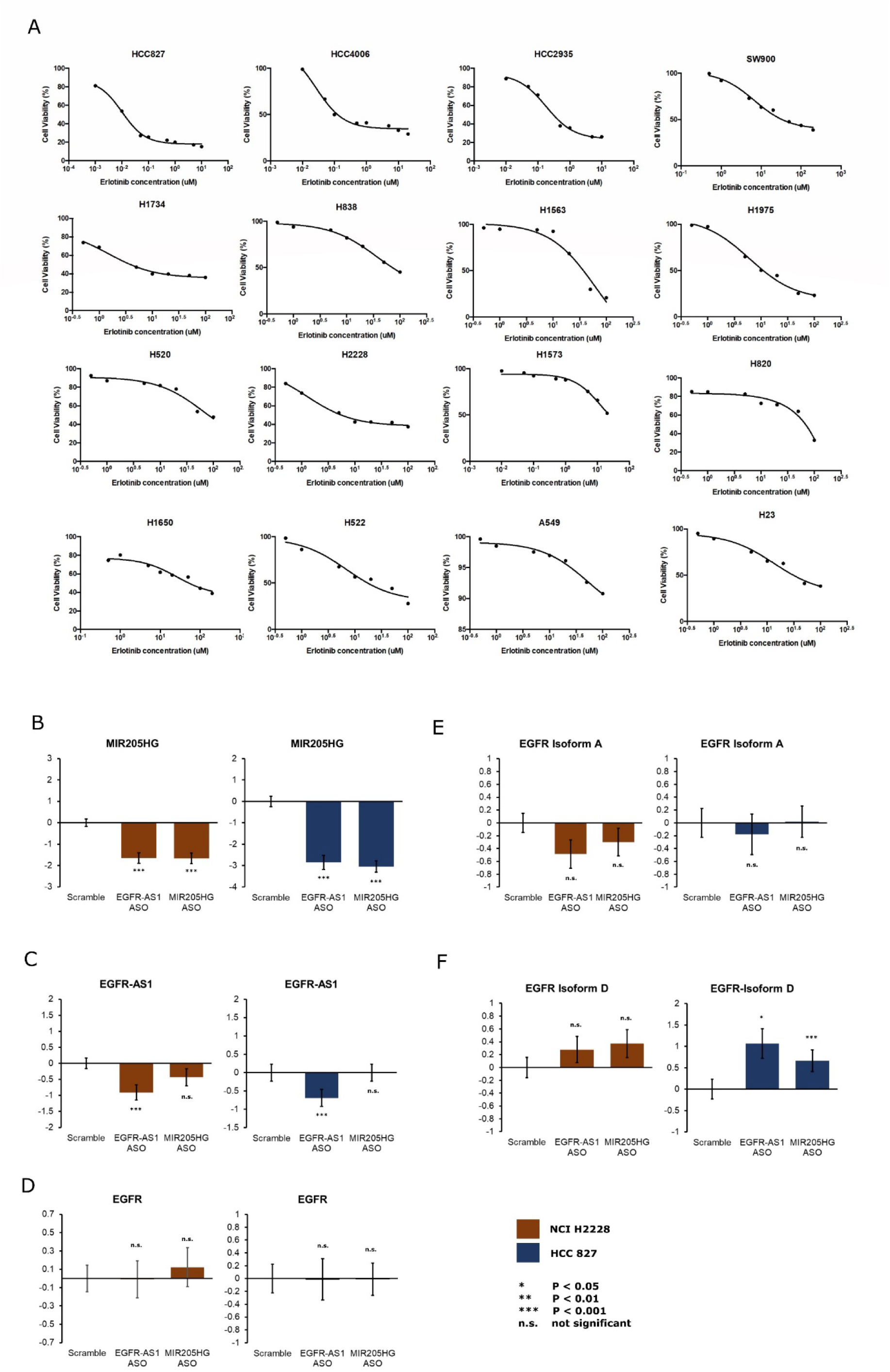
Dose-response and qPCR experiments in lung cancer cell lines. **A.** Dose-response curves of erlotinib treatment in 16 lung cancer cell lines. **B-F.** Expression levels of *MIR205HG* (B), *EGFR-AS1* (C), *EGFR* (D), *EGFR* Isoform A (E), *EGFR* Isoform D (F) determined using qRT-PCR in NCI H2228 and HCC 827 cell lines with ASO-mediated k.d. of *EGFR-AS1* and *MIR205HG*. The mean and t-test p-values were obtained from six independent biological replicates

Supplementary Table 1: ANOVA and Tukey’s post-hoc analysis of lncRNA biotype distribution in GDSC and CTRP gene expression datasets

Supplementary Table 2: LASSO coefficients of drug response modeled using PCG-lncRNA co-expression modules, GO and KEGG enrichment analysis of modules

Supplementary Table 3: Drug response regression coefficients (effects) and p-values associated with oncogenic and tumor suppressor lncRNAs

